## Supplementary Information for "Principles for enhancing virus capsid capacity and stability from a thermophilic virus capsid structure"

### Slide 1
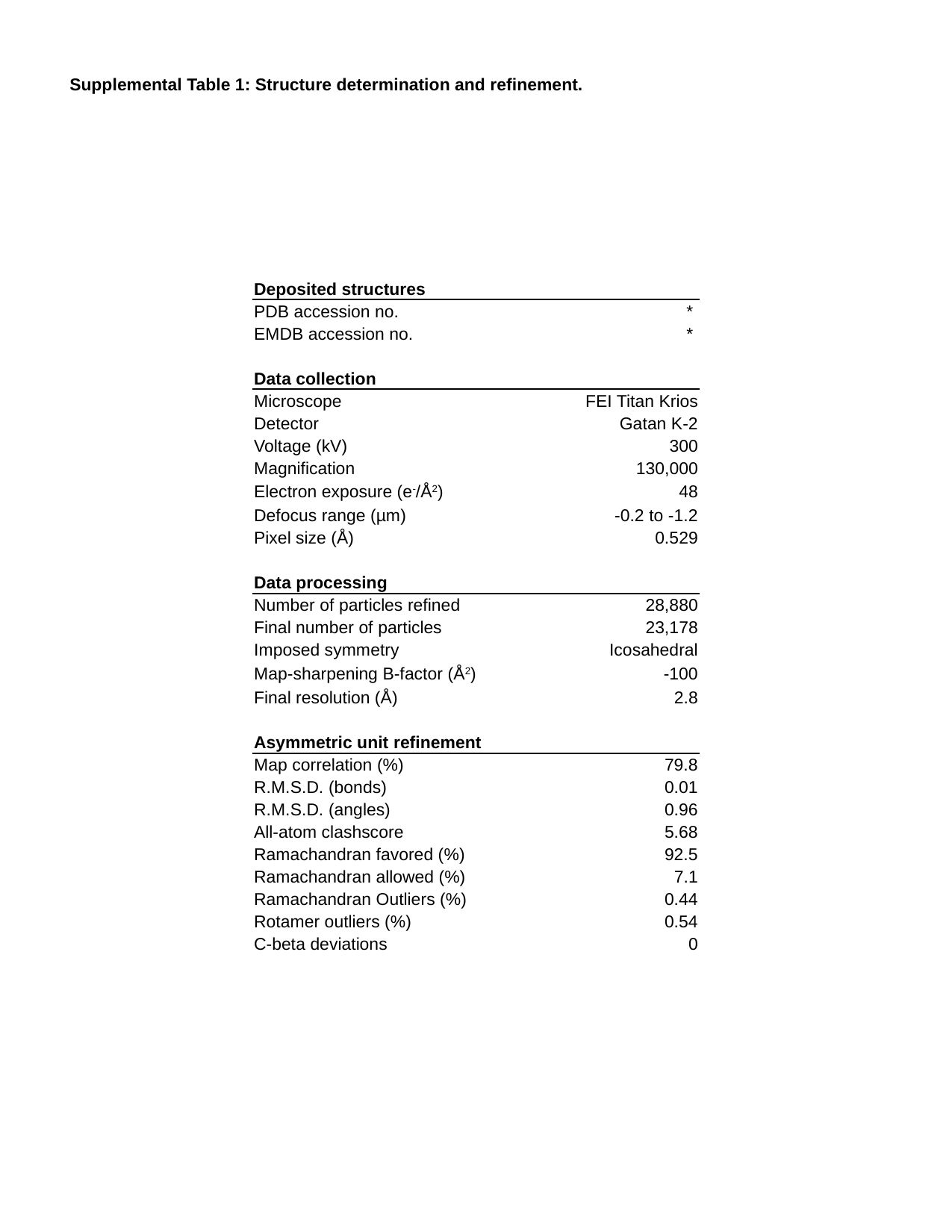

Supplemental Table 1: Structure determination and refinement.
| Deposited structures | |
| --- | --- |
| PDB accession no. | \* |
| EMDB accession no. | \* |
| Data collection | |
| Microscope | FEI Titan Krios |
| Detector | Gatan K-2 |
| Voltage (kV) | 300 |
| Magnification | 130,000 |
| Electron exposure (e^-^/Å^2^) | 48 |
| Defocus range (µm) | -0.2 to -1.2 |
| Pixel size (Å) | 0.529 |
| Data processing | |
| Number of particles refined | 28,880 |
| Final number of particles | 23,178 |
| Imposed symmetry | Icosahedral |
| Map-sharpening B-factor (Å^2^) | -100 |
| Final resolution (Å) | 2.8 |
| Asymmetric unit refinement | |
| Map correlation (%) | 79.8 |
| R.M.S.D. (bonds) | 0.01 |
| R.M.S.D. (angles) | 0.96 |
| All-atom clashscore | 5.68 |
| Ramachandran favored (%) | 92.5 |
| Ramachandran allowed (%) | 7.1 |
| Ramachandran Outliers (%) | 0.44 |
| Rotamer outliers (%) | 0.54 |
| C-beta deviations | 0 |

### Slide 2
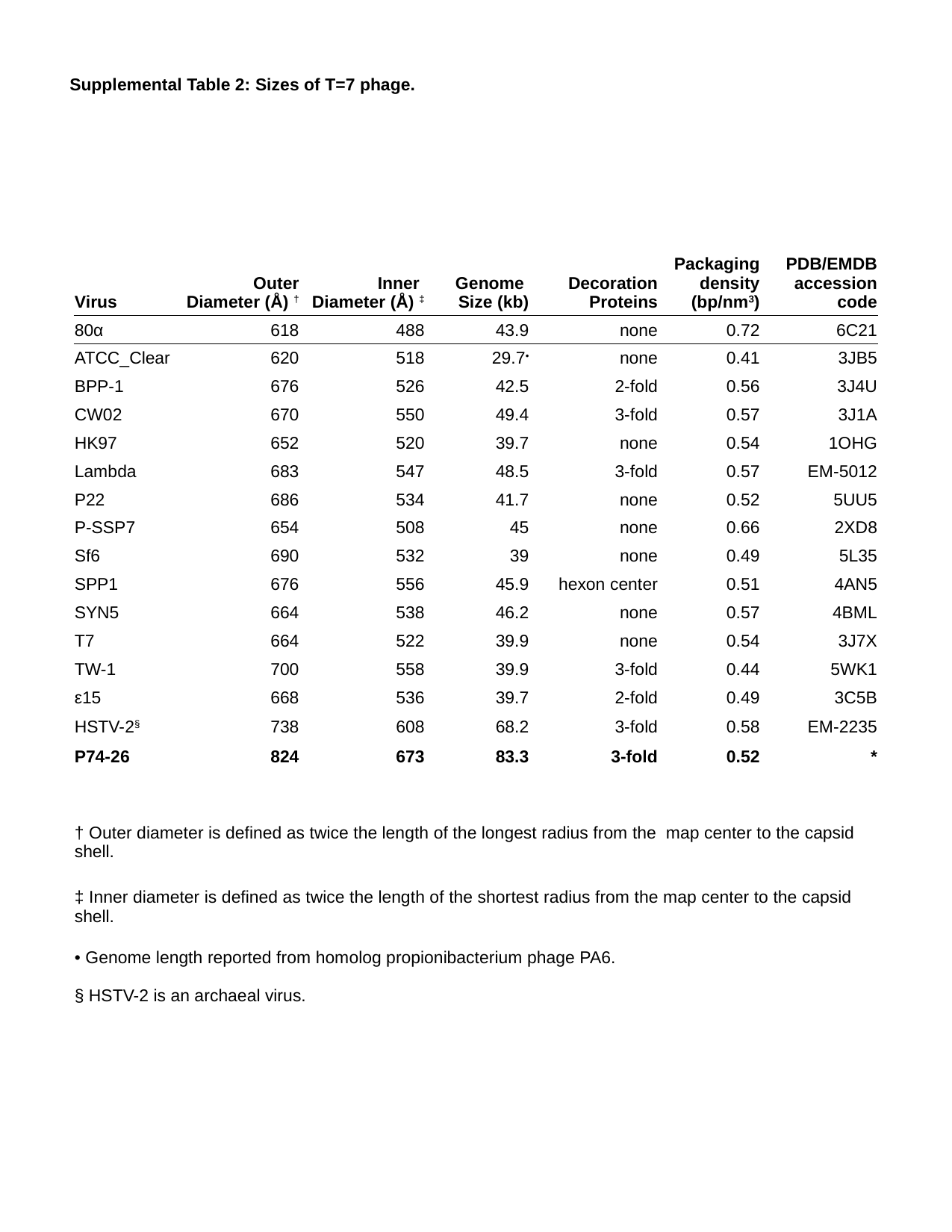

Supplemental Table 2: Sizes of T=7 phage.
| Virus | Outer Diameter (Å) ^†^ | Inner Diameter (Å) ^‡^ | Genome Size (kb) | Decoration Proteins | Packaging density (bp/nm^3^) | PDB/EMDB accession code |
| --- | --- | --- | --- | --- | --- | --- |
| 80α | 618 | 488 | 43.9 | none | 0.72 | 6C21 |
| ATCC\_Clear | 620 | 518 | 29.7^•^ | none | 0.41 | 3JB5 |
| BPP-1 | 676 | 526 | 42.5 | 2-fold | 0.56 | 3J4U |
| CW02 | 670 | 550 | 49.4 | 3-fold | 0.57 | 3J1A |
| HK97 | 652 | 520 | 39.7 | none | 0.54 | 1OHG |
| Lambda | 683 | 547 | 48.5 | 3-fold | 0.57 | EM-5012 |
| P22 | 686 | 534 | 41.7 | none | 0.52 | 5UU5 |
| P-SSP7 | 654 | 508 | 45 | none | 0.66 | 2XD8 |
| Sf6 | 690 | 532 | 39 | none | 0.49 | 5L35 |
| SPP1 | 676 | 556 | 45.9 | hexon center | 0.51 | 4AN5 |
| SYN5 | 664 | 538 | 46.2 | none | 0.57 | 4BML |
| T7 | 664 | 522 | 39.9 | none | 0.54 | 3J7X |
| TW-1 | 700 | 558 | 39.9 | 3-fold | 0.44 | 5WK1 |
| ε15 | 668 | 536 | 39.7 | 2-fold | 0.49 | 3C5B |
| HSTV-2^§^ | 738 | 608 | 68.2 | 3-fold | 0.58 | EM-2235 |
| P74-26 | 824 | 673 | 83.3 | 3-fold | 0.52 | \* |
| † Outer diameter is defined as twice the length of the longest radius from the map center to the capsid shell. |
| --- |
| ‡ Inner diameter is defined as twice the length of the shortest radius from the map center to the capsid shell. |
| • Genome length reported from homolog propionibacterium phage PA6. § HSTV-2 is an archaeal virus. |

### Slide 3
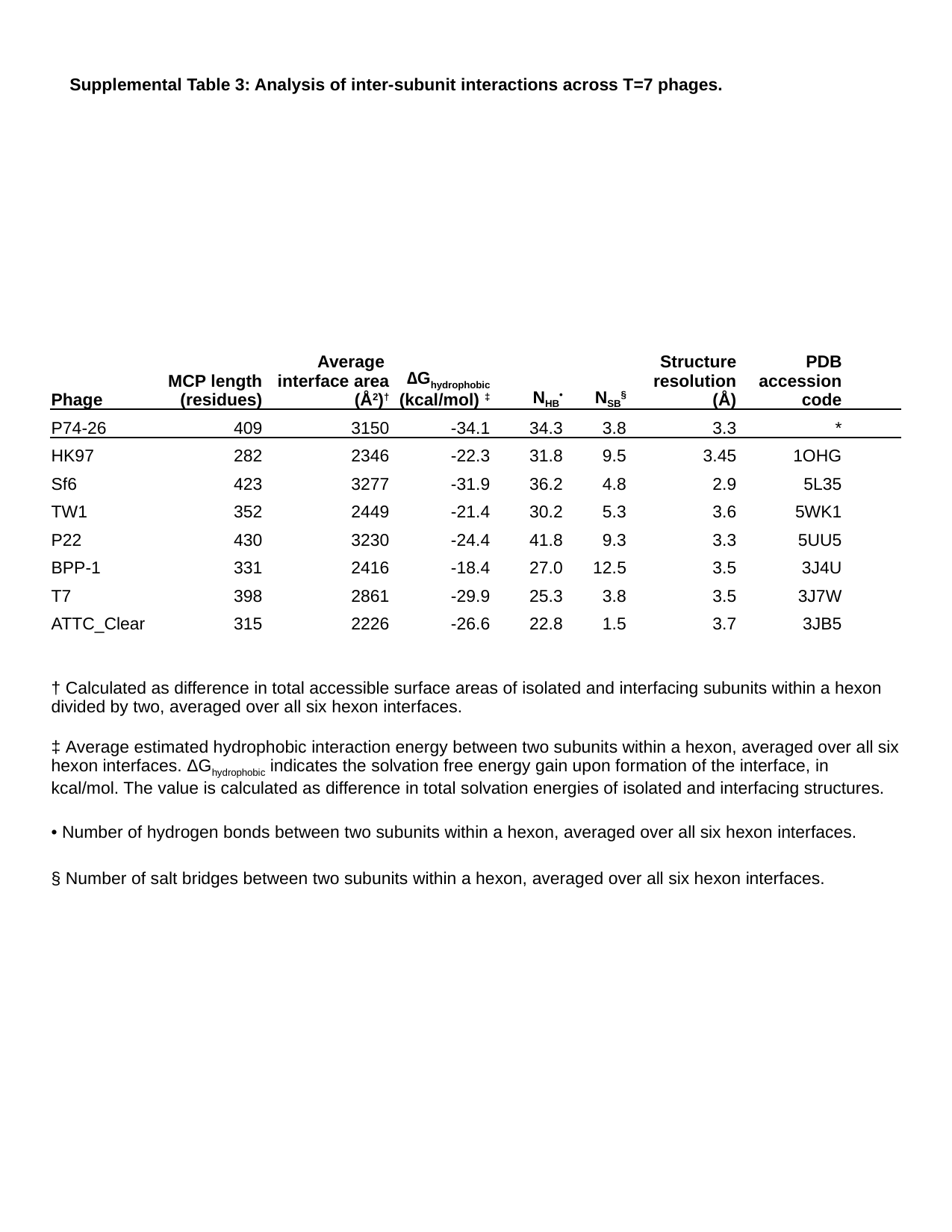

Supplemental Table 3: Analysis of inter-subunit interactions across T=7 phages.
| Phage | MCP length (residues) | Average interface area (Å^2^)^†^ | ∆G_hydrophobic_ (kcal/mol) ^‡^ | N_HB_^•^ | N_SB_^§^ | Structure resolution (Å) | PDB accession code |
| --- | --- | --- | --- | --- | --- | --- | --- |
| P74-26 | 409 | 3150 | -34.1 | 34.3 | 3.8 | 3.3 | \* |
| HK97 | 282 | 2346 | -22.3 | 31.8 | 9.5 | 3.45 | 1OHG |
| Sf6 | 423 | 3277 | -31.9 | 36.2 | 4.8 | 2.9 | 5L35 |
| TW1 | 352 | 2449 | -21.4 | 30.2 | 5.3 | 3.6 | 5WK1 |
| P22 | 430 | 3230 | -24.4 | 41.8 | 9.3 | 3.3 | 5UU5 |
| BPP-1 | 331 | 2416 | -18.4 | 27.0 | 12.5 | 3.5 | 3J4U |
| T7 | 398 | 2861 | -29.9 | 25.3 | 3.8 | 3.5 | 3J7W |
| ATTC\_Clear | 315 | 2226 | -26.6 | 22.8 | 1.5 | 3.7 | 3JB5 |
| † Calculated as difference in total accessible surface areas of isolated and interfacing subunits within a hexon divided by two, averaged over all six hexon interfaces. | | | | | | | |
| ‡ Average estimated hydrophobic interaction energy between two subunits within a hexon, averaged over all six hexon interfaces. ΔG_hydrophobic_ indicates the solvation free energy gain upon formation of the interface, in kcal/mol. The value is calculated as difference in total solvation energies of isolated and interfacing structures. | | | | | | | |
| • Number of hydrogen bonds between two subunits within a hexon, averaged over all six hexon interfaces. | | | | | | | |
| § Number of salt bridges between two subunits within a hexon, averaged over all six hexon interfaces. | | | | | | | |

### Slide 4
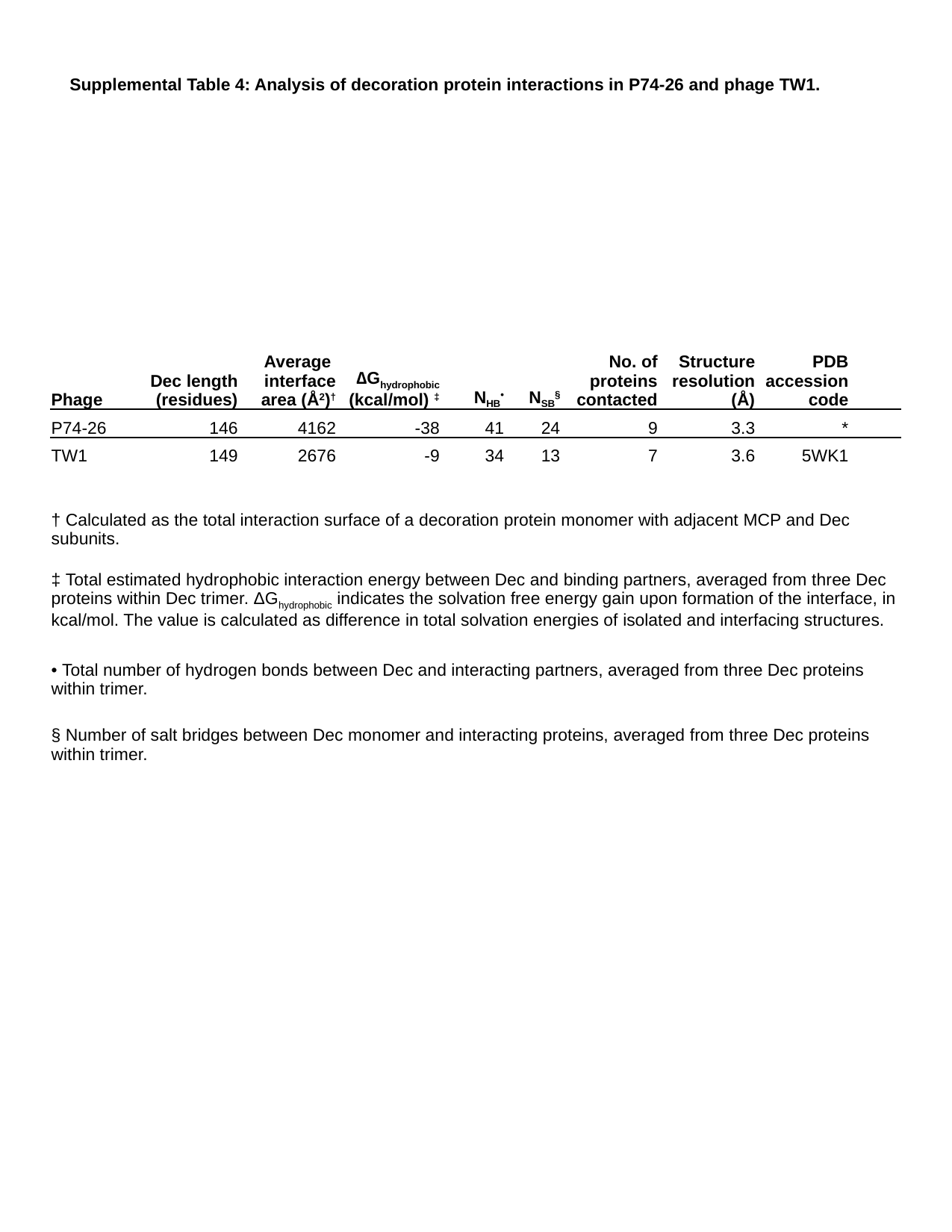

Supplemental Table 4: Analysis of decoration protein interactions in P74-26 and phage TW1.
| Phage | Dec length (residues) | Average interface area (Å^2^)^†^ | ∆G_hydrophobic_ (kcal/mol) ^‡^ | N_HB_^•^ | N_SB_^§^ | No. of proteins contacted | Structure resolution (Å) | PDB accession code |
| --- | --- | --- | --- | --- | --- | --- | --- | --- |
| P74-26 | 146 | 4162 | -38 | 41 | 24 | 9 | 3.3 | \* |
| TW1 | 149 | 2676 | -9 | 34 | 13 | 7 | 3.6 | 5WK1 |
| † Calculated as the total interaction surface of a decoration protein monomer with adjacent MCP and Dec subunits. | | | | | | | | |
| ‡ Total estimated hydrophobic interaction energy between Dec and binding partners, averaged from three Dec proteins within Dec trimer. ΔG_hydrophobic_ indicates the solvation free energy gain upon formation of the interface, in kcal/mol. The value is calculated as difference in total solvation energies of isolated and interfacing structures. | | | | | | | | |
| • Total number of hydrogen bonds between Dec and interacting partners, averaged from three Dec proteins within trimer. | | | | | | | | |
| § Number of salt bridges between Dec monomer and interacting proteins, averaged from three Dec proteins within trimer. | | | | | | | | |

### Slide 5
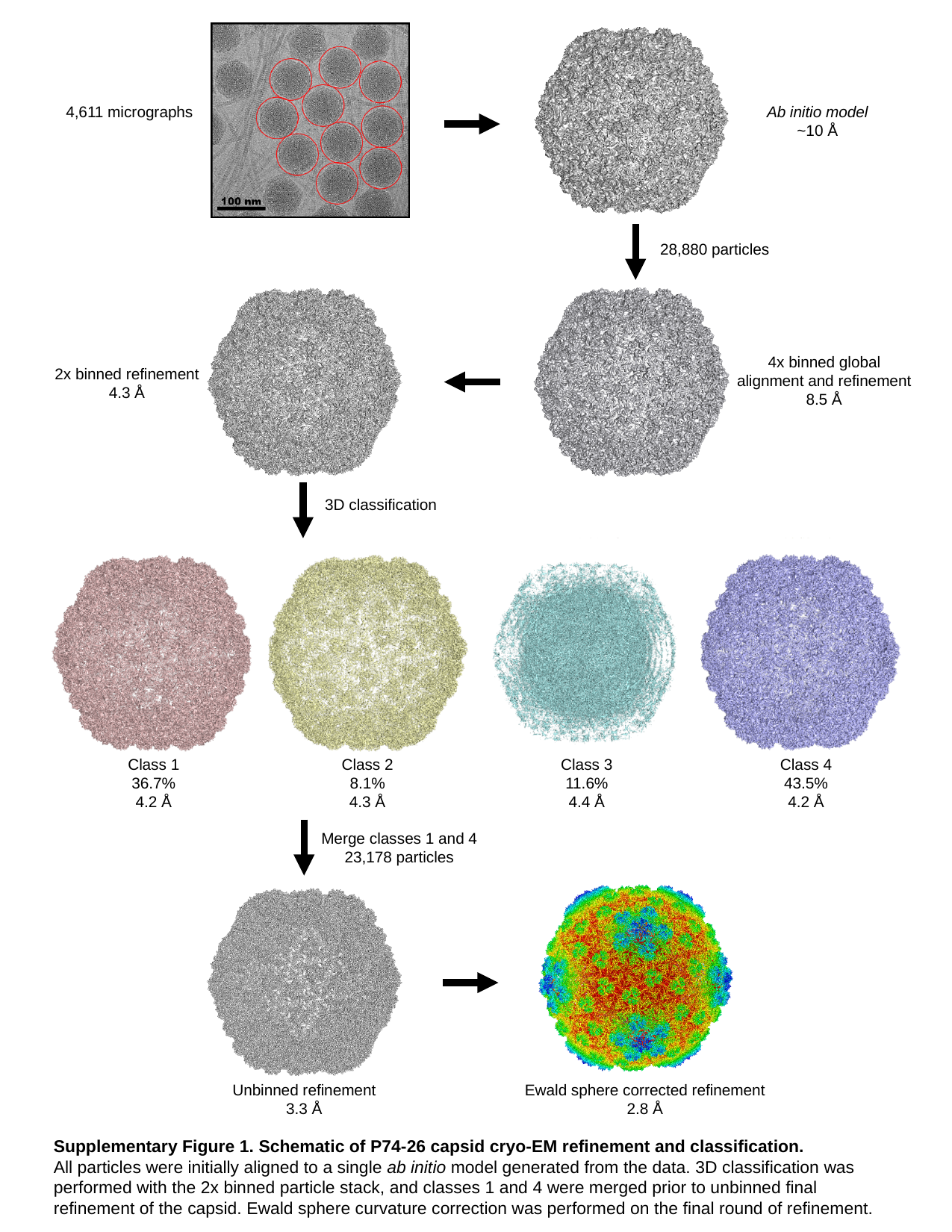

4,611 micrographs
Ab initio model
~10 Å
28,880 particles
4x binned global alignment and refinement
8.5 Å
2x binned refinement
4.3 Å
Class 3
11.6%
4.4 Å
Class 4
43.5%
4.2 Å
Class 1
36.7%
4.2 Å
Class 2
8.1%
4.3 Å
3D classification
Merge classes 1 and 4
23,178 particles
Unbinned refinement
3.3 Å
Ewald sphere corrected refinement
2.8 Å
Supplementary Figure 1. Schematic of P74-26 capsid cryo-EM refinement and classification.
All particles were initially aligned to a single ab initio model generated from the data. 3D classification was performed with the 2x binned particle stack, and classes 1 and 4 were merged prior to unbinned final refinement of the capsid. Ewald sphere curvature correction was performed on the final round of refinement.

### Slide 6
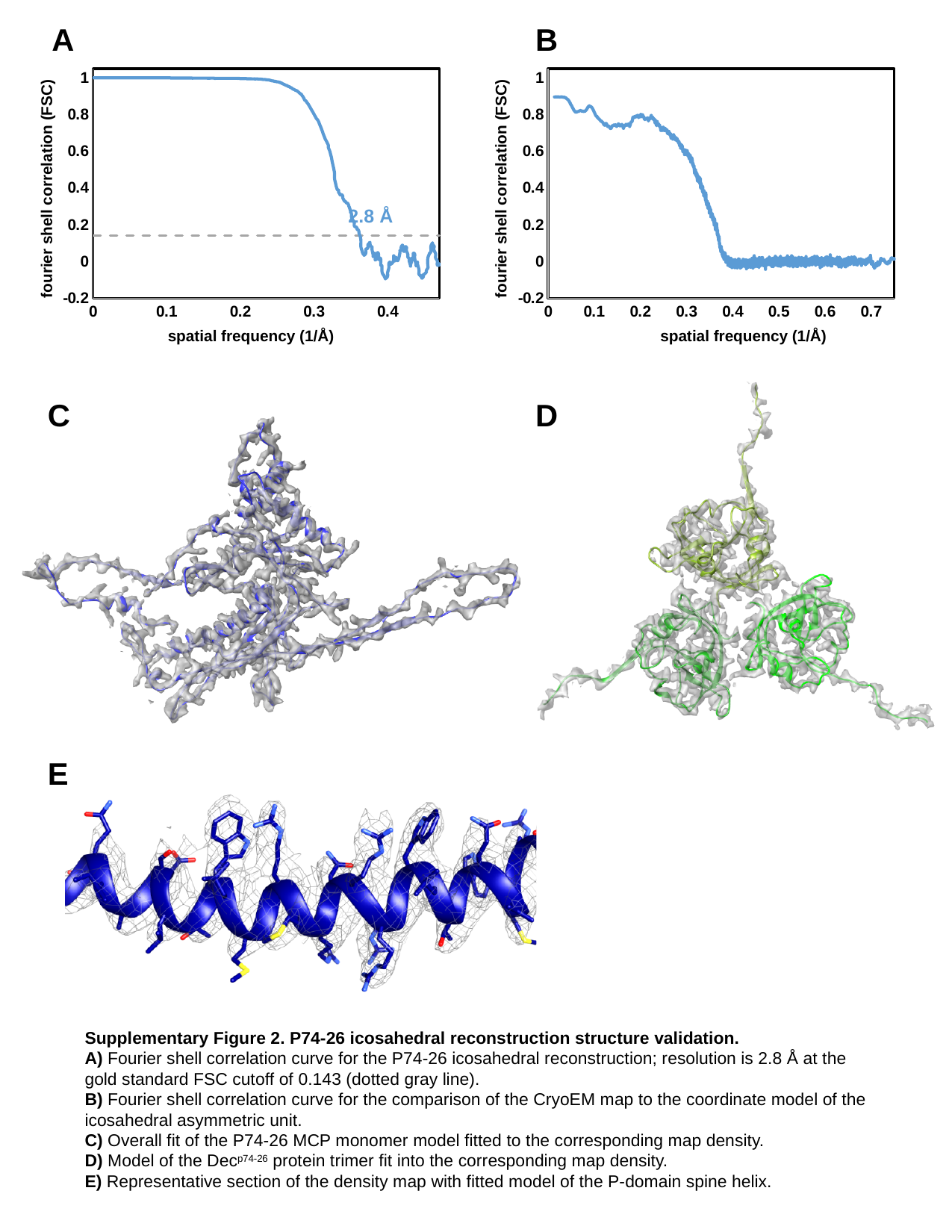

B
A
#### Chart
| Category | y | y |
|---|---|---|
#### Chart
| Category | y |
|---|---|fourier shell correlation (FSC)
fourier shell correlation (FSC)
2.8 Å
spatial frequency (1/Å)
spatial frequency (1/Å)
C
D
E
Supplementary Figure 2. P74-26 icosahedral reconstruction structure validation.
A) Fourier shell correlation curve for the P74-26 icosahedral reconstruction; resolution is 2.8 Å at the gold standard FSC cutoff of 0.143 (dotted gray line).
B) Fourier shell correlation curve for the comparison of the CryoEM map to the coordinate model of the icosahedral asymmetric unit.
C) Overall fit of the P74-26 MCP monomer model fitted to the corresponding map density.
D) Model of the Dec^p74-26^ protein trimer fit into the corresponding map density.
E) Representative section of the density map with fitted model of the P-domain spine helix.

### Slide 7
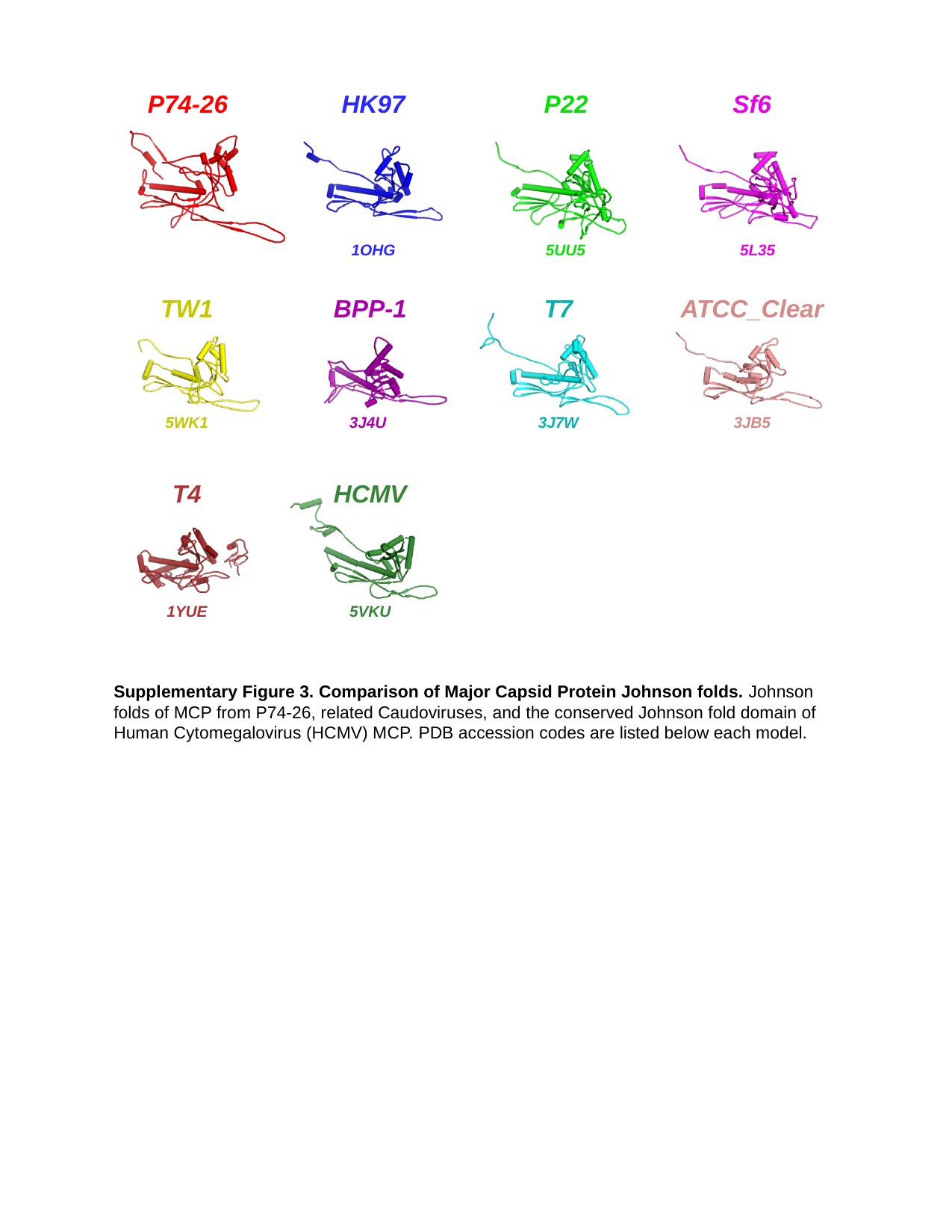

P74-26
HK97
P22
Sf6
1OHG
5UU5
5L35
TW1
BPP-1
ATCC_Clear
T7
3J7W
3JB5
3J4U
5WK1
T4
HCMV
1YUE
5VKU
Supplementary Figure 3. Comparison of Major Capsid Protein Johnson folds. Johnson folds of MCP from P74-26, related Caudoviruses, and the conserved Johnson fold domain of Human Cytomegalovirus (HCMV) MCP. PDB accession codes are listed below each model.

### Slide 8
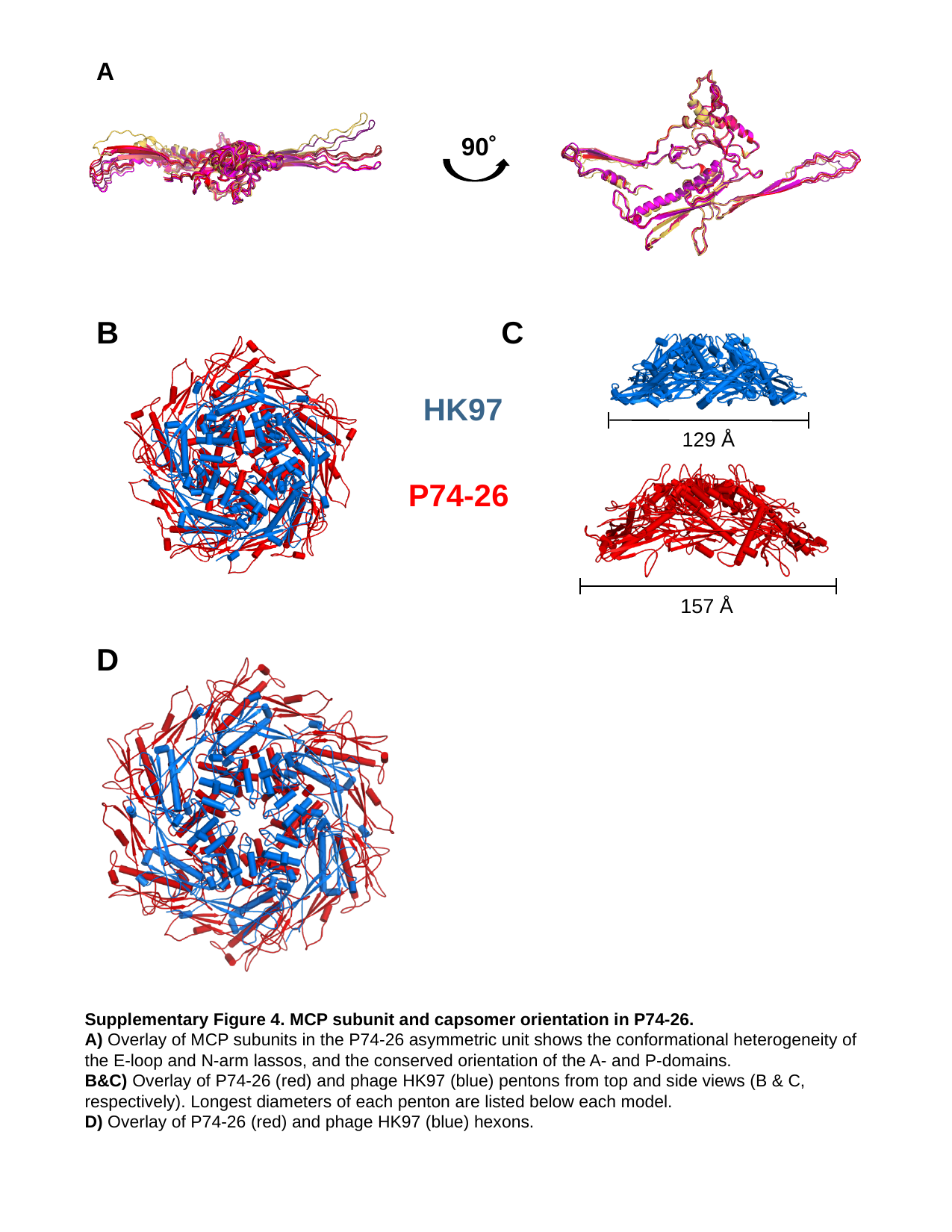

A
90˚
B
C
HK97
129 Å
P74-26
157 Å
D
Supplementary Figure 4. MCP subunit and capsomer orientation in P74-26.
A) Overlay of MCP subunits in the P74-26 asymmetric unit shows the conformational heterogeneity of the E-loop and N-arm lassos, and the conserved orientation of the A- and P-domains.
B&C) Overlay of P74-26 (red) and phage HK97 (blue) pentons from top and side views (B & C, respectively). Longest diameters of each penton are listed below each model.
D) Overlay of P74-26 (red) and phage HK97 (blue) hexons.

### Slide 9
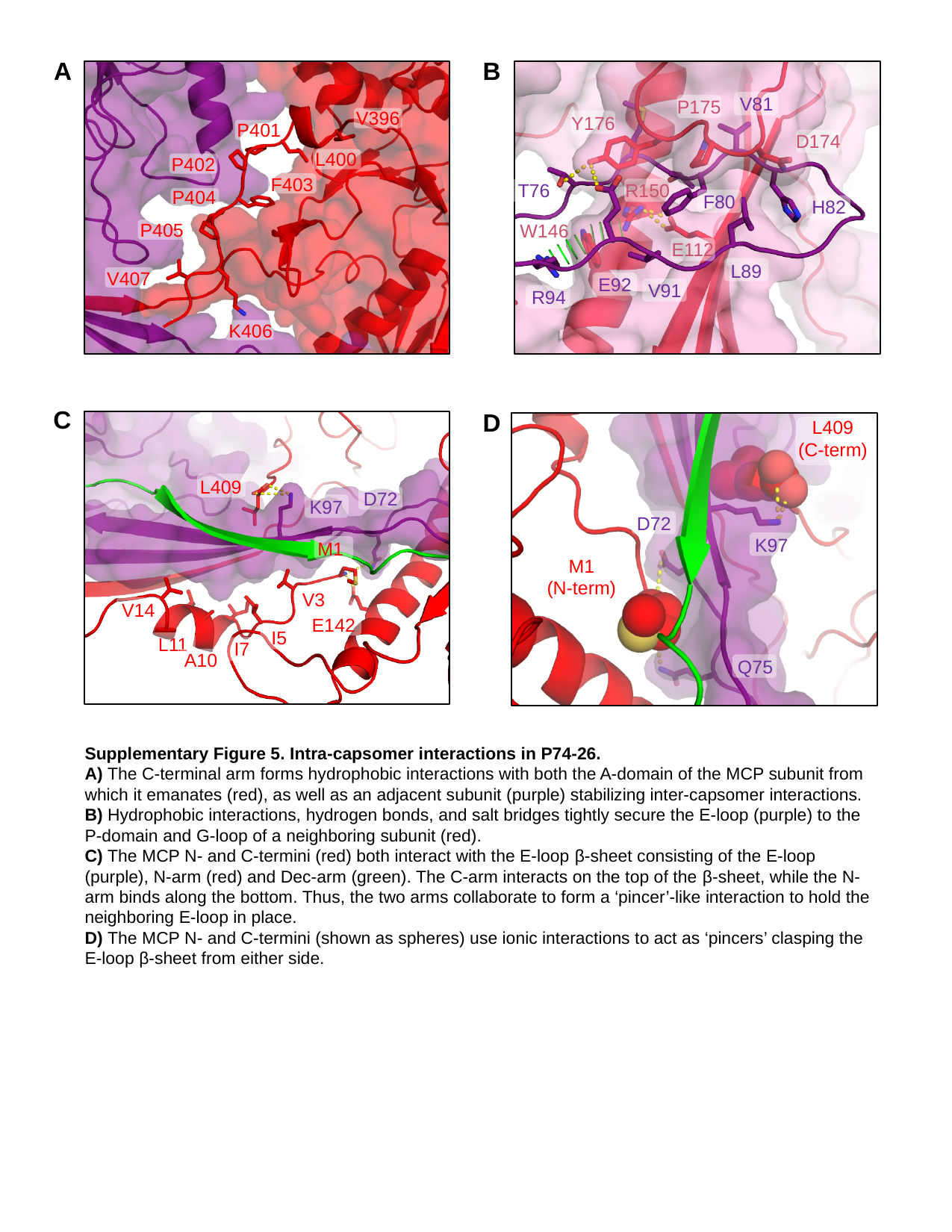

A
B
V396
P401
L400
P402
F403
P404
P405
V407
K406
V81
P175
Y176
D174
T76
R150
F80
H82
W146
E112
L89
E92
V91
R94
C
L409
D72
K97
M1
V3
V14
E142
I5
L11
I7
A10
D
L409
(C-term)
D72
K97
M1
(N-term)
Q75
Supplementary Figure 5. Intra-capsomer interactions in P74-26.
A) The C-terminal arm forms hydrophobic interactions with both the A-domain of the MCP subunit from which it emanates (red), as well as an adjacent subunit (purple) stabilizing inter-capsomer interactions.
B) Hydrophobic interactions, hydrogen bonds, and salt bridges tightly secure the E-loop (purple) to the P-domain and G-loop of a neighboring subunit (red).
C) The MCP N- and C-termini (red) both interact with the E-loop β-sheet consisting of the E-loop (purple), N-arm (red) and Dec-arm (green). The C-arm interacts on the top of the β-sheet, while the N-arm binds along the bottom. Thus, the two arms collaborate to form a ‘pincer’-like interaction to hold the neighboring E-loop in place.
D) The MCP N- and C-termini (shown as spheres) use ionic interactions to act as ‘pincers’ clasping the E-loop β-sheet from either side.

### Slide 10
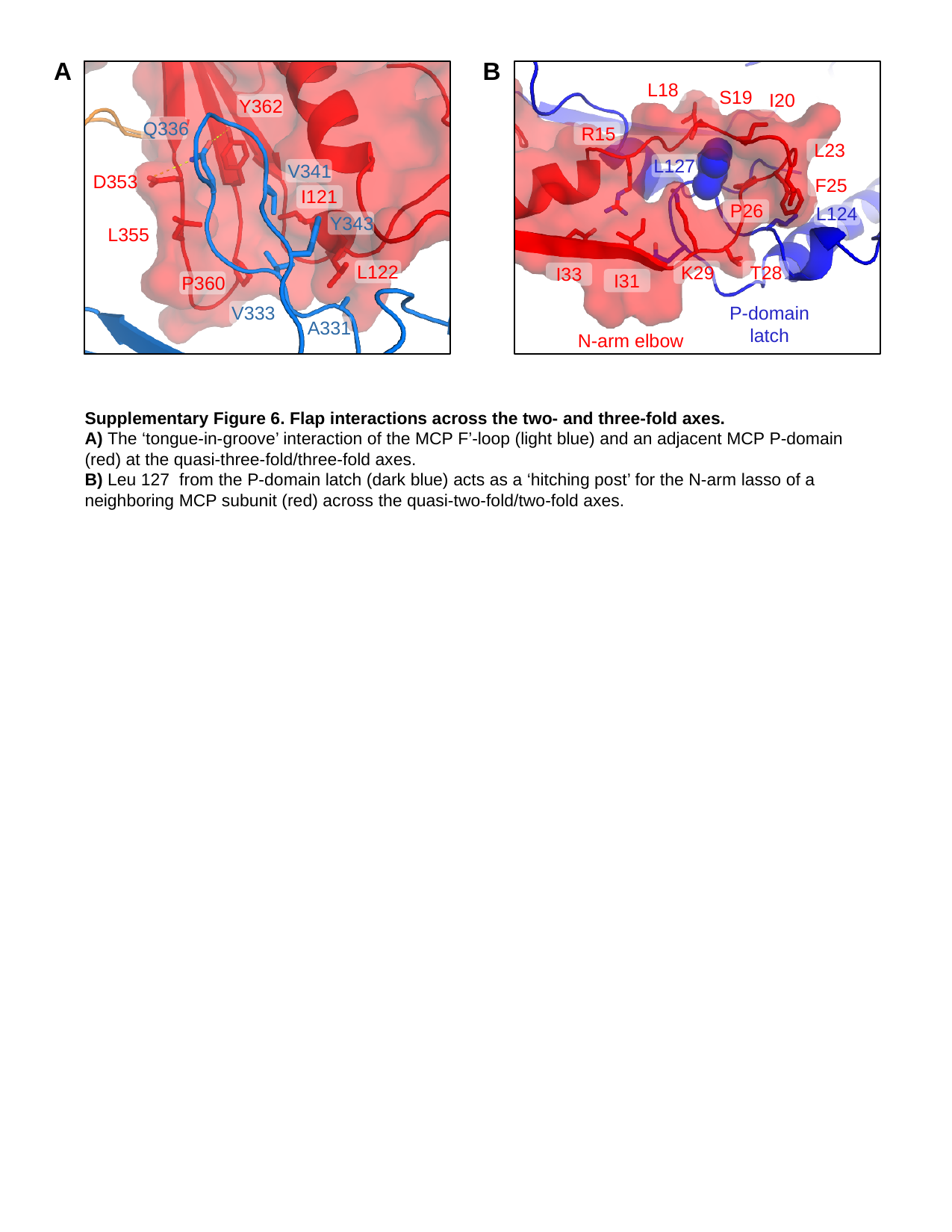

A
B
V396
P401
L400
P402
F403
P404
P405
V407
K406
L18
S19
I20
Y362
Q336
R15
L23
L127
V341
D353
F25
I121
P26
L124
Y343
L355
L122
K29
T28
I33
I31
P360
V333
P-domain
latch
A331
N-arm elbow
Supplementary Figure 6. Flap interactions across the two- and three-fold axes.
A) The ‘tongue-in-groove’ interaction of the MCP F’-loop (light blue) and an adjacent MCP P-domain (red) at the quasi-three-fold/three-fold axes.
B) Leu 127 from the P-domain latch (dark blue) acts as a ‘hitching post’ for the N-arm lasso of a neighboring MCP subunit (red) across the quasi-two-fold/two-fold axes.

### Slide 11
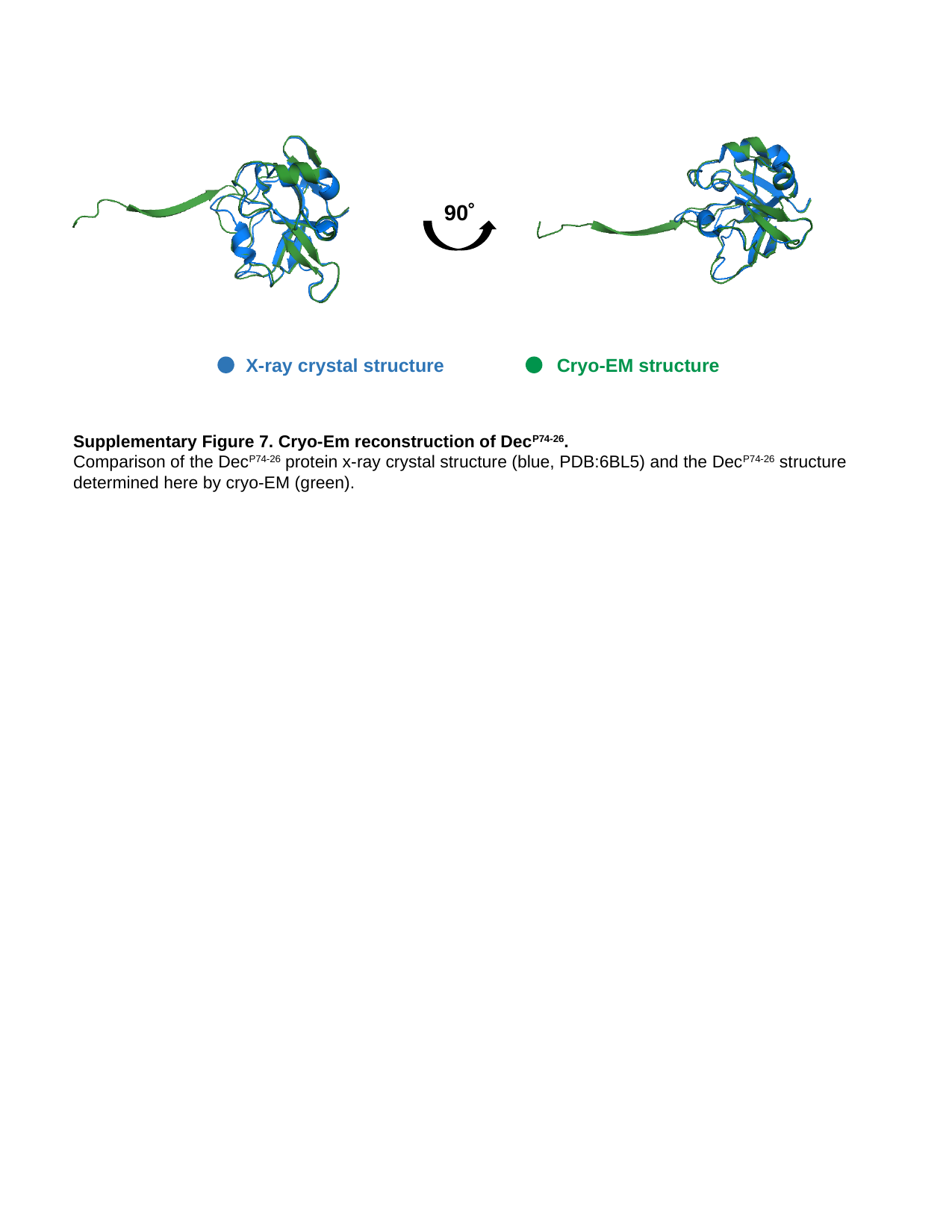

90˚
X-ray crystal structure
Cryo-EM structure
Supplementary Figure 7. Cryo-Em reconstruction of Dec^P74-26^.
Comparison of the Dec^P74-26^ protein x-ray crystal structure (blue, PDB:6BL5) and the Dec^P74-26^ structure determined here by cryo-EM (green).

### Slide 12
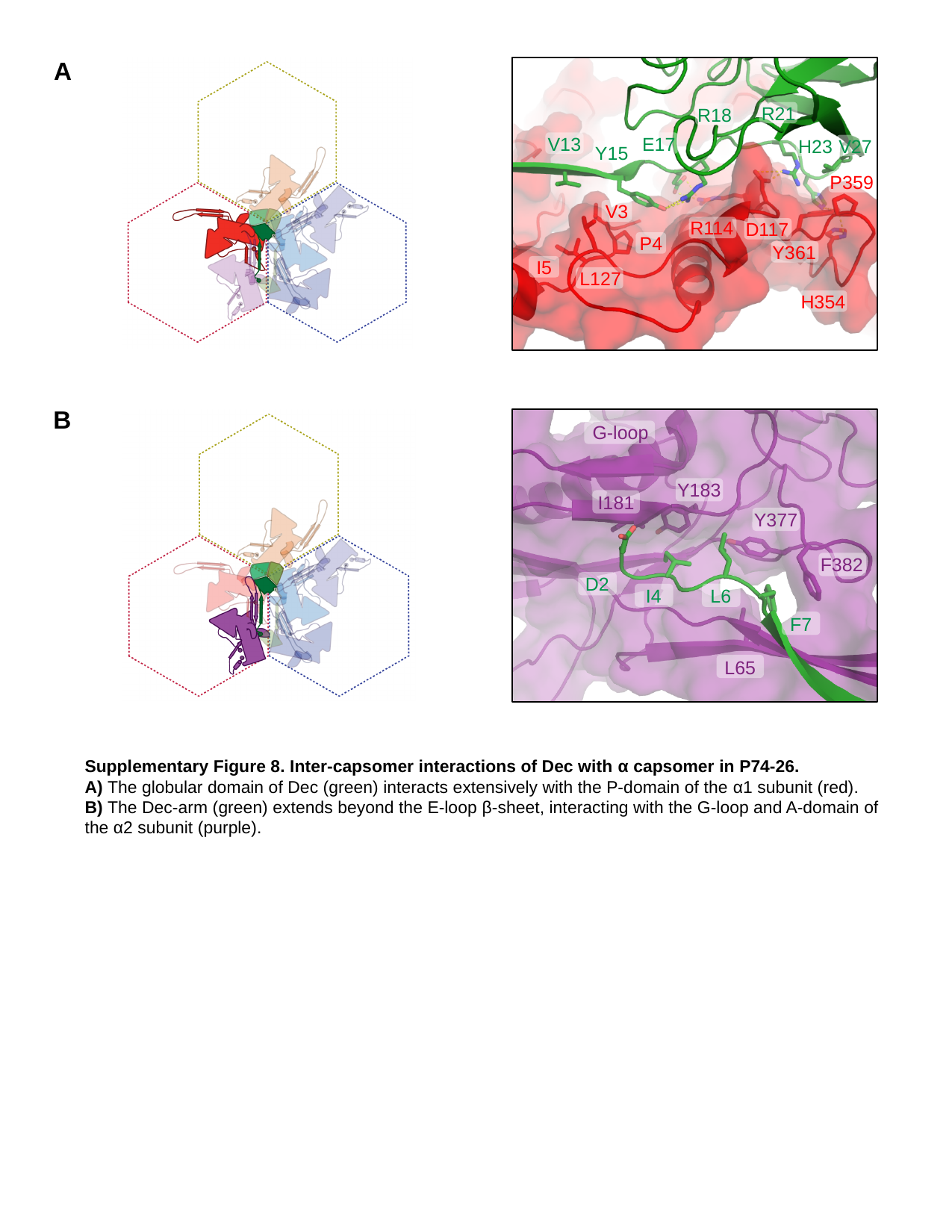

A
R21
R18
V13
E17
H23
V27
Y15
P359
V3
R114
D117
P4
Y361
I5
L127
H354
B
G-loop
Y183
I181
Y377
F382
D2
I4
L6
F7
L65
Supplementary Figure 8. Inter-capsomer interactions of Dec with α capsomer in P74-26.
A) The globular domain of Dec (green) interacts extensively with the P-domain of the α1 subunit (red).
B) The Dec-arm (green) extends beyond the E-loop β-sheet, interacting with the G-loop and A-domain of the α2 subunit (purple).

### Slide 13
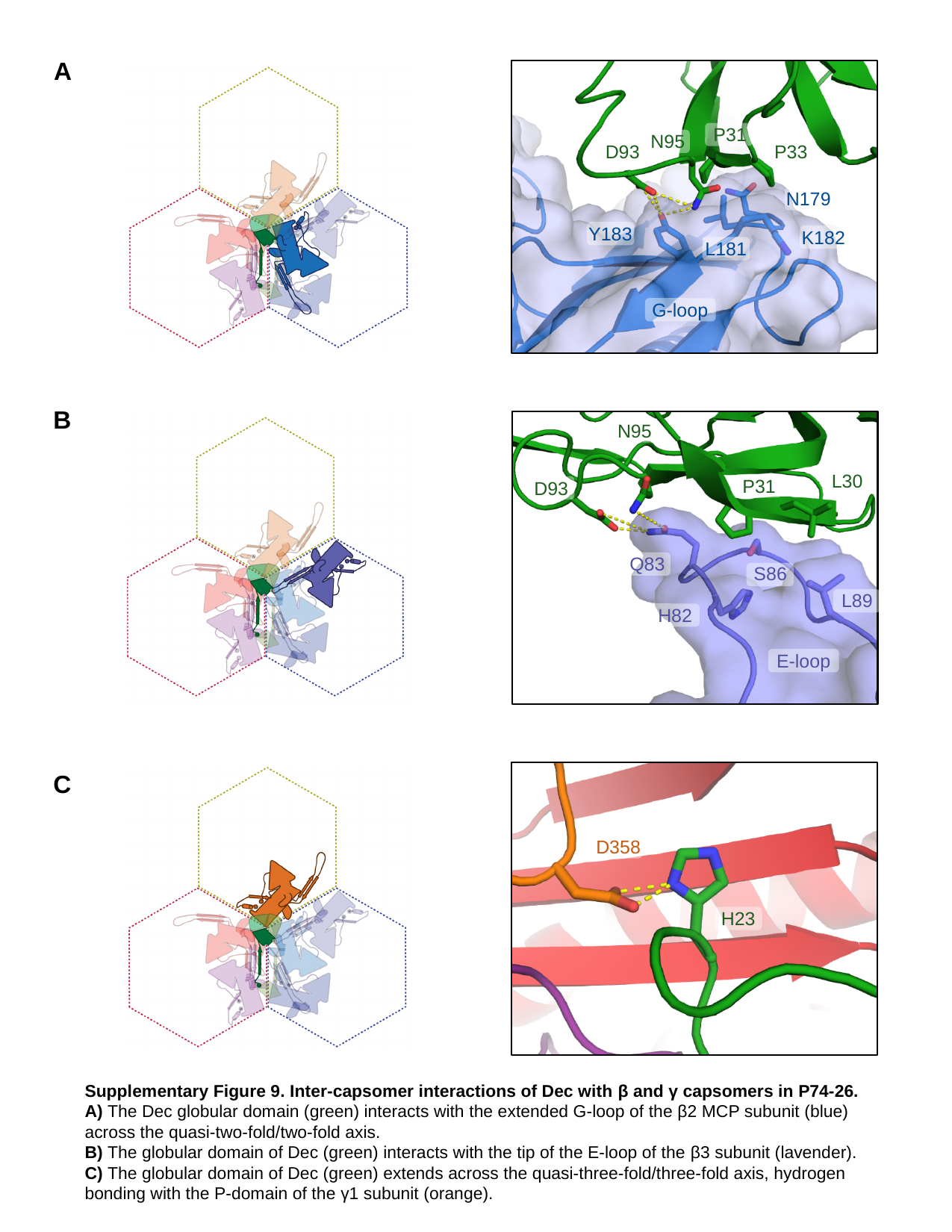

A
P31
N95
D93
P33
N179
Y183
K182
L181
G-loop
B
N95
L30
P31
D93
Q83
S86
L89
H82
E-loop
C
D358
H23
Supplementary Figure 9. Inter-capsomer interactions of Dec with β and γ capsomers in P74-26.
A) The Dec globular domain (green) interacts with the extended G-loop of the β2 MCP subunit (blue) across the quasi-two-fold/two-fold axis.
B) The globular domain of Dec (green) interacts with the tip of the E-loop of the β3 subunit (lavender).
C) The globular domain of Dec (green) extends across the quasi-three-fold/three-fold axis, hydrogen bonding with the P-domain of the γ1 subunit (orange).

### Slide 14
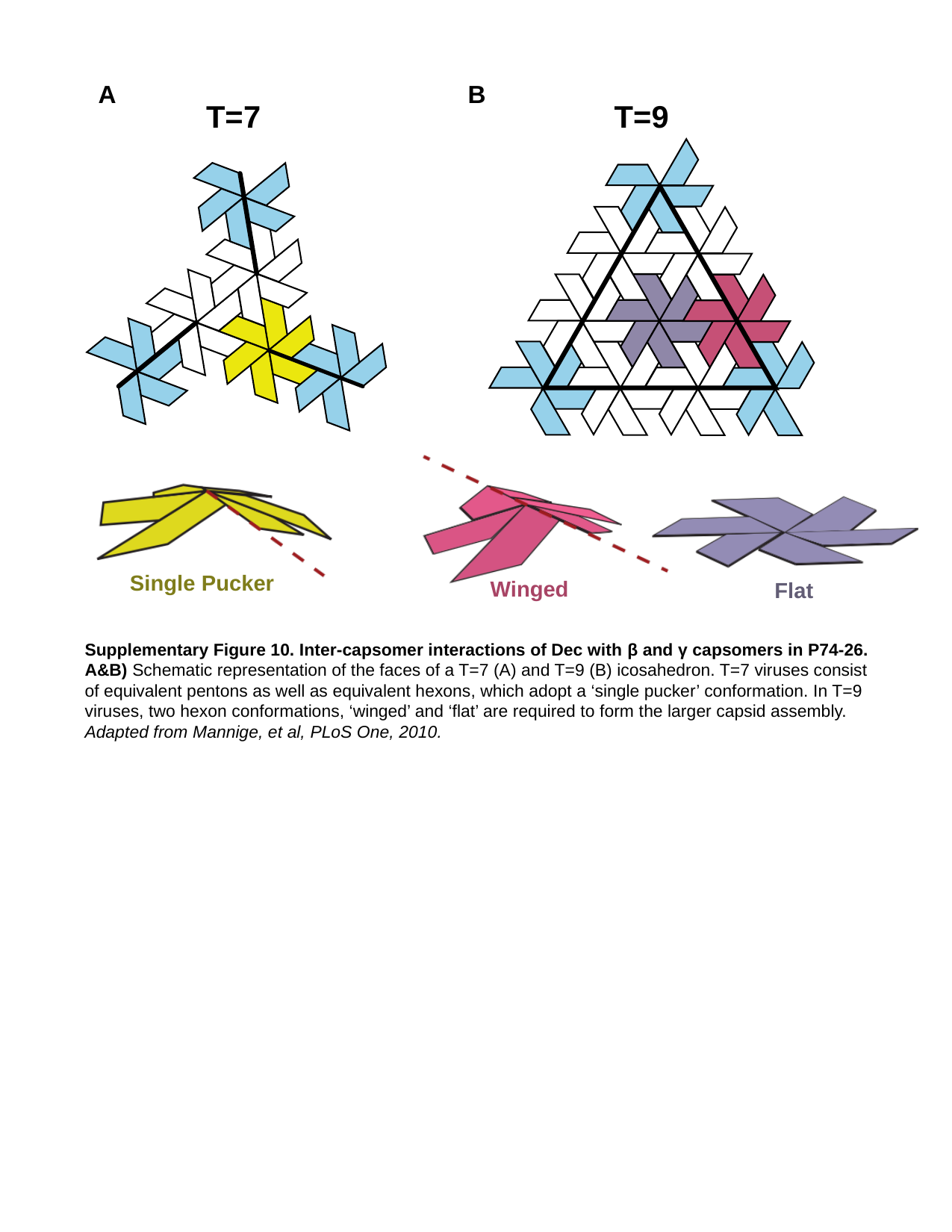

A
B
T=9
T=7
Single Pucker
Winged
Flat
Supplementary Figure 10. Inter-capsomer interactions of Dec with β and γ capsomers in P74-26.
A&B) Schematic representation of the faces of a T=7 (A) and T=9 (B) icosahedron. T=7 viruses consist of equivalent pentons as well as equivalent hexons, which adopt a ‘single pucker’ conformation. In T=9 viruses, two hexon conformations, ‘winged’ and ‘flat’ are required to form the larger capsid assembly. Adapted from Mannige, et al, PLoS One, 2010.
